# Mash Screen: High-throughput sequence containment estimation for genome discovery

**DOI:** 10.1101/557314

**Authors:** Brian D Ondov, Gabriel J Starrett, Anna Sappington, Aleksandra Kostic, Sergey Koren, Christopher B Buck, Adam M Phillippy

**Affiliations:** Genome Informatics Section, National Human Genome Research Institute, Bethesda, MD, USA; Department of Computer Science, University of Maryland College Park, College Park, MD, USA; Tumor Virus Molecular Biology Section, National Cancer Institute, Bethesda, MD, USA; Department of Electrical Engineering and Computer Science, Massachusetts Institute of Technology, Cambridge, MA, USA; Department of Computer Science, Princeton University, Princeton, NJ, USA

**Keywords:** sample, article, author

## Abstract

The MinHash algorithm has proven effective for rapidly estimating the resemblance of two genomes or metagenomes. However, this method cannot reliably estimate the containment of a genome within a metagenome. Here we describe an online algorithm capable of measuring the containment of genomes and proteomes within either assembled or unassembled sequencing read sets. We describe several use cases, including contamination screening and retrospective analysis of metagenomes for novel genome discovery. Using this tool, we provide containment estimates for every NCBI RefSeq genome within every SRA metagenome, and demonstrate the identification of a novel polyomavirus species from a public metagenome.

## 1 Introduction

As evolving sequencing technology continues to increase throughput and lower costs, databases of sequenced genomes (e.g. NCBI RefSeq [1]) continue their exponential growth, making searches against them ever more complex [2, 3]. Furthermore, the body of raw sequencing data, NCBI’s Sequence Read Archive (SRA) [4], is growing even faster, outpacing our ability to assemble and curate the genomes and metagenomes represented [5, 6]. These trends have popularized alignment-free analytical methods, typically operating on k-mers, which are short (∼21 bp), overlapping subsequences of the genome for some length *k*. Our prior work in this area introduced Mash [7], which uses the MinHash dimensionality-reduction technique [8] to compress k-mer sets of whole genomes to *sketches* of just several hundred or thousand values, enabling extremely fast estimations of genomic or metagenomic distances. Though useful for isolated genomes or high-level metagenomic correlation, Mash is ill-suited for analyzing the constituents of a metagenomic mixture. This is because typical applications of MinHashing approximate *resemblance* rather than *containment*. In this work we address the containment problem and present a new method, Mash Screen, for measuring how well a genome or proteome is represented within a metagenome. This has applications ranging from quick contamination screens to the tracking of specific bacterial and viral strains across all available metagenomes.

Though originally defined in terms of documents and words, here we will consider the resemblance and containment of biological sequences and their constituent k-mers. For two genomes *a* and *b*, their resemblance is defined as the Jaccard index *j* between their k-mer sets *A* and *B*,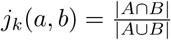. Thus, resemblance is sensitive to differential sequence lengths. Consider a single genome *a* and a much larger metagenome *b*. The resemblance between *a* and *b* is expected to be low, even if *b* wholly contains *a*, because *b* will likely contain many k-mers not present in *a*, making the union of *A* and *B* larger than the intersection. Resemblance ranges from 0 to 1, is symmetric, and 1 *- j* is a metric. In contrast, the containment index *c* is asymmetric and measures the extent to which one genome is contained in another, 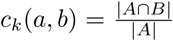. If all the k-mers of *A* are contained in *B*, regardless of how many other k-mers *B* contains, the containment index will equal 1. Because genome sequences can be very long, we seek to efficiently approximate *j*_*k*_(*a, b*) and *c*_*k*_(*a, b*) by considering only a small fraction of the total k-mers.

For estimating resemblance, Mash uses a ‘bottom sketch’ strategy as originally proposed by Broder [8]. More efficient techniques for estimating resemblance have since emerged [9, 10, 11, 12], but bottom sketching is elegant in its simplicity. In short, all k-mers from a genome *A* are passed through a single hash function *h* but only the smallest *m* hash values are stored as the sketch *S*(*A*), where |*S*(*A*)| *<<* |*A*|. Because the probability that *S*(*A*) and *S*(*B*) share a minimum value is related to the Jaccard index of *A* and *B*, the resemblance of the full k-mer sets *A* and *B* can be quickly approximated from the comparatively smaller sketches *S*(*A*) and *S*(*B*). Mash further converts resemblance to the Mash distance, which is an estimate of mutational distance between the two sequences.

While the difference in formulation between resemblance and containment is small, different strategies are required to accurately approximate them. In contrast to resemblance, which can be reliably estimated using fixed-size sketches, estimating containment requires a sketch size proportional to the genome size (Fig. 1). To accomplish this, Broder originally proposed using a modulo function for containment sketching, which produces proportionally-sized sketches. However, the modulo sketch has the disadvantage of requiring a very large sketch size in order to achieve acceptable error bounds when searching for small contained elements, such as viruses. To bound the size of this data structure, Bloom filters have been suggested as a replacement for modulus sketches when estimating containment [13]. In the Bloom filter scheme, |*A∩B*| is estimated using a MinHash sketch of *A* and a Bloom filter of *B*. Alternatively, Bloom trees [14] and Bloom filter matrices [15] have been proposed for the general task of detecting smaller sequences within large sequence read collections. However, such indexes have only been applicable transcriptomes or isolate genomes, rather than metagenomes.

**Figure 1.**
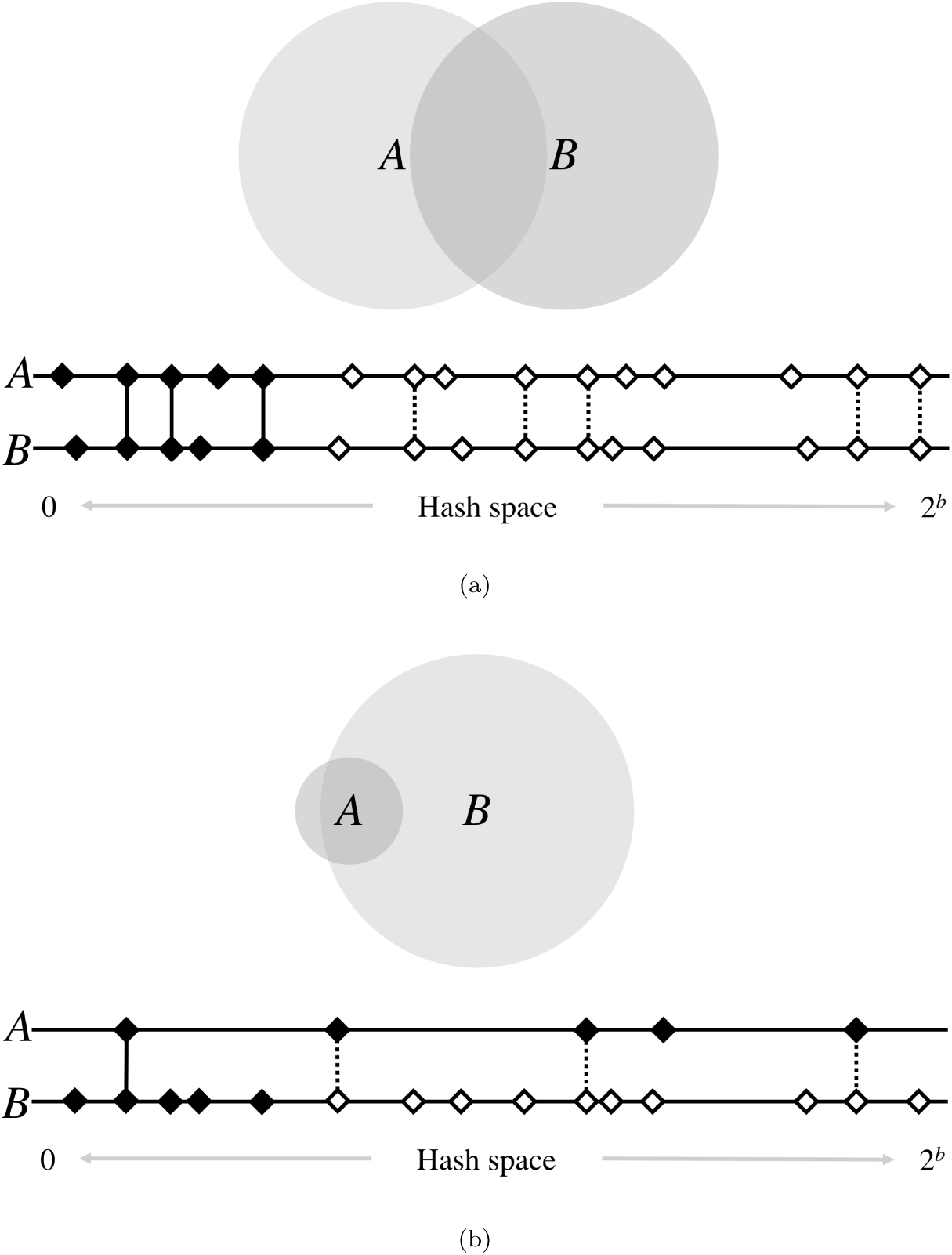
High-level diagram of set operations with MinHash. The size of the shaded circles represents k-mer set sizes, while the area of their overlap represents the cardinality of their intersections. Below each pair of circles is a diagram of the MinHash resemblance estimation for a sketch size of 5. Horizontal lines represent the space of possible hash values, of which there are 2^*b*^, where *b* is the number of bits used for hashing. Here, diamonds are hashes of the k-mers in sets *A* and *B*, and black shading indicates the smallest 5 hashes in each. Vertical lines indicate matching hashes and are solid only if both hashes are in the bottom sketch of their respective set. In (a), genomes of similar sizes are well-suited for resemblance estimation, since their hashes are similarly distributed across the hash space. Matching hashes are usually both in, or both out, of the bottom 5. However, if the genomes are of very different sizes, as in (b), the larger genome will saturate the space more densely. This causes a higher fraction of matching hashes to be contained in only the sketch of the smaller set, underestimating the containment of *A* in *B*. Thus, all hashes of *B* must be considered to accurately estimate the containment of *A*.

These prior containment estimation methods have all sought to compress and/or index a large collection of sequencing reads, with the goal of enabling rapid sequence search across them (e.g. given a reference database of metagenomes, identify all those containing some query sequence). This is an important problem, but in this paper we consider the converse: given a reference database of genomes, identify all those contained in some query metagenome. We would also like to estimate, without assembly, the similarity between the reference genomes and those contained in the metagenome.

To address this problem, we propose the concept of a *screen*, in which we rapidly test a database of many genomes for their containment within a larger, unassembled metagenome (Fig. 2). For each reference genome, Mash Screen computes a containment score that measures the similarity of the reference genome to that which is contained within the metagenome. This enables routine tasks such as quick contamination screening; the selection of appropriate reference genomes for mapping-based analyses; and efficient screening of entire metagenomic databases for tracking individual species and strains across all samples. For the latter case, we have computed the containment of every NCBI RefSeq genome within every NCBI SRA metagenome, and have made these results available for download and query. To demonstrate the potential utility of this dataset, we describe the identification and assembly of a novel polyomavirus species from a public SRA metagenome.

**Figure 2.**
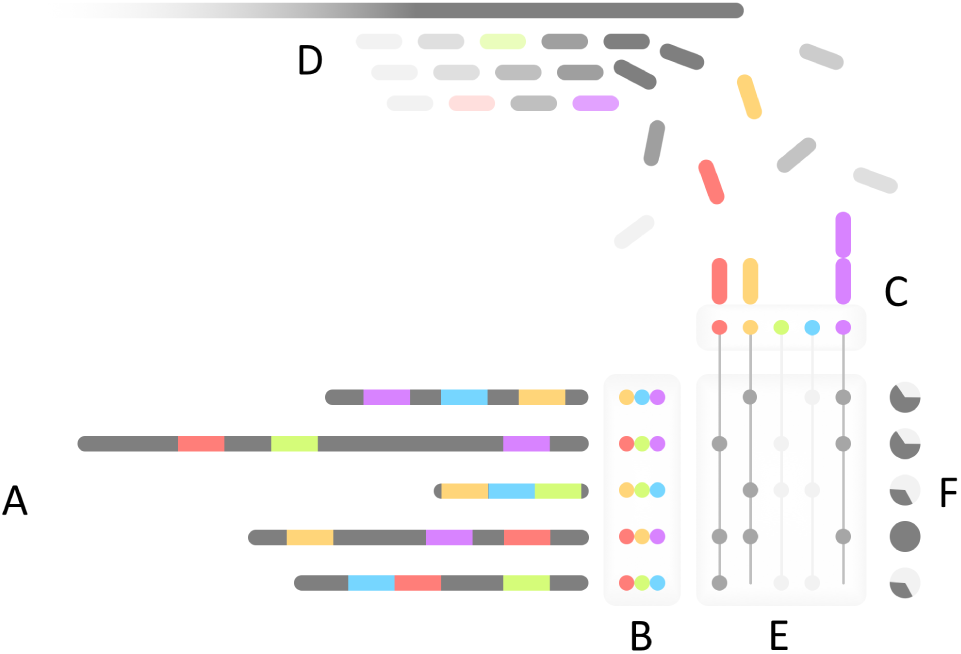
Mash Screen algorithmic overview. (A) The minimum *m* hashes (in this case 3, shown colored) for each reference sequence is determined during sketching to produce (B) a reference MinHash sketch library. For screening, distinct hashes from all reference sketches are collected and used as keys to (C) a map of observed counts per hash, which is populated by (D) hashing k-mers from the sequence mixture as it is streamed. (E) Counts from the map are queried for each sketch to produce (F) a containment estimation for each constituent of the mixture.

## 2 Results

### 2.1 Correlation between Mash distance and containment estimates

For the k-mers of a given reference genome *A*, a related genome *A*^*′*^ that is contained within a metagenomic read set *B* should, by design, have a Mash containment score *C*_*k*_(*A, B*) that is inversely correlated with the Mash distance *D*_*k*_(*A, A*^*′*^). Ideally, *C*_*k*_(*A, B*) = 1 *- D*(*A, A*^*′*^). However, because the metagenome may contain k-mers matching *A* that are not from *A*^*′*^, and the different formulations of the Mash distance and containment, the correlation is imperfect.

To test how well Mash distance correlates with containment, we analyzed metagenomic reads, both real and simulated, from a mock community of bacteria and archaea with known constituents [16], referred to here as the Shakya dataset. All RefSeq genomes were sketched, and those genomes sharing at least one sketch value with a known Shakya constituent were collected, resulting in 82,086 genomes. Wholegenome Mash distances between each constituent genome and the RefSeq subset were then compared to Mash containment scores for each RefSeq genome versus the metagenome. Screening the 11.1 gigabases of Illumina reads against RefSeq for this analysis took 61 CPU minutes.

Mash containment scores showed good correlation for Mash distances less than 0.2 (*r*^2^= 0.99), but tended to overestimate Mash distances greater than 0.2 (*r*^2^= 0.54 for all points, Fig. 3a). First, like Mash distance estimates, the expected error of the containment estimates increases for more diverged sequences. Second, when estimating containment, k-mers from any genome in the mixture are permitted to match the reference, whereas the Mash distance is restricted to matching only k-mers from the genome being compared. Figure 3 (left) also highlights a number of outliers that sit well off the diagonal. Points circled in red represent RefSeq genomes that do not share high similarity to the expected constituent genomes, yet have high containment scores when compared to the metagenome. This indicates suspected contamination in the Shakya mock community.

**Figure 3.**
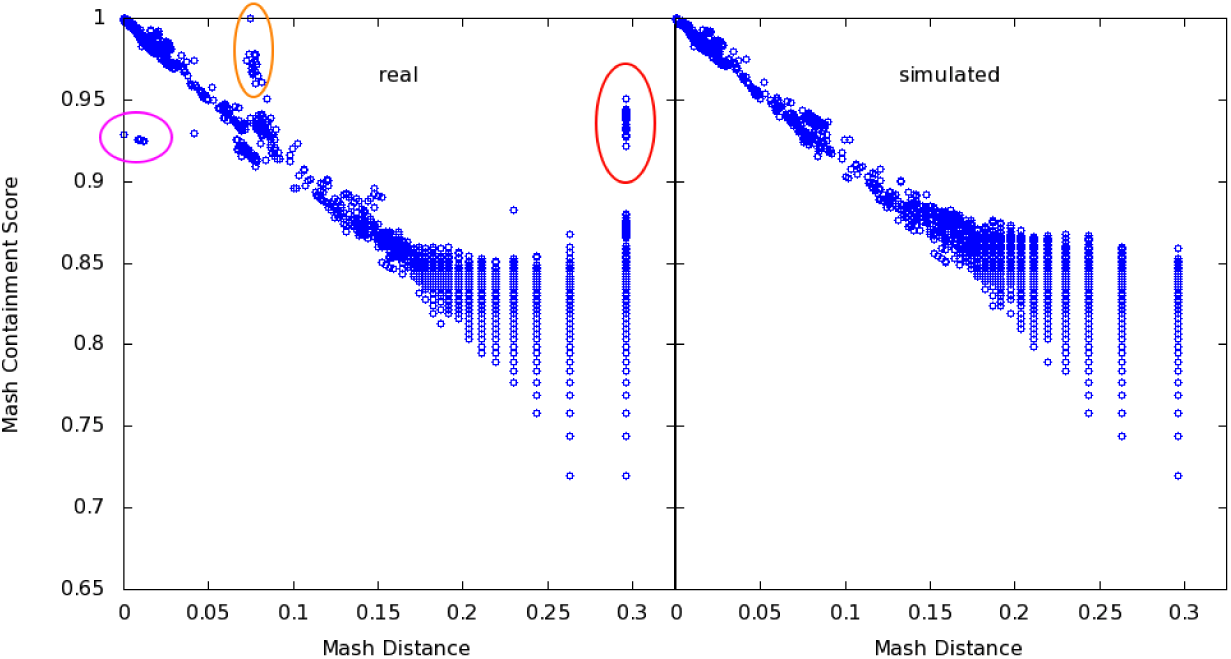
Correlation of Mash containment scores with pairwise Mash distances. Points represent RefSeq genomes with at least one matching hash to the known reference constituents of the Shakya mock microbial community, which serves as an expected distance (*x*-axis). Raw reads from this metagenome were run through Mash Screen to obtain a containment score for each RefSeq genome (*y*-axis). Left, real Illumina reads sequenced from the mock community (SRR606249). The area circled in red highlights RefSeq genomes with higher than expected containment scores caused by contamination of the sequencing run. This contamination was independently confirmed via read mapping, revealing the presence of at least four additional species. Circled in magenta and orange are two clades of *F. nucleatum* that are consistent with low abundance of the reference strain (magenta) and intra-species contamination (orange), both of which have been previously described for this dataset. Right, reads simulated from only the known constituents corrects the outliers and yields the expected correlation.

*E.coli* and *P. ruminis* contamination had been previously detected in this mock community by Awad *et al.* [17], and so we speculated that this was causing the discrepancy between Mash distance and containment estimates. Consistent with this explanation, simulating reads from only the listed constituents improved overall correlation to *r*^2^= 0.73 (Fig. 3, right). Further investigation of the outlier points revealed additional *P. acnes* and *S. parasanguinis* contamination that was not identified by the prior studies. Mapping all metagenomic reads to these reference genomes confirmed the presence of the four contaminating species or their near relatives (Table 1). After filtering points arising from these contaminants, correlation of the real dataset was identical to that of the simulated dataset (*r*^2^= 0.73). Also notable are points closer to the diagonal, but still in distinct clusters, highlighted in magenta and orange in Figure 3 (left). These represent two distinct clades of *nucleatum*, a gram-negative commensal of the human oral cavity that has well-delineated subspecies [18]. The expected reference strain, ATCC 25586, was noted by Awad *et al.* to be present in low abundance, which is likely what is causing the Mash containment to be lower than expected for this and similar strains (below the diagonal, circled in magenta). The strains above the diagonal (circled in orange), however, represent *F. nucleatum* subspecies (*polymorphum* and *vincentii*) that Awad *et al.* suggested could be intraspecies contaminants. The results of this experiment are thus widely in concordance with prior, independent research, lending validation to both the past findings and to our new method.

**Table 1.** Identified contaminants in the Shakya dataset measured by mapped reads and Mash Screen. *Organism* refers to the closest strain in RefSeq to the suspected contaminant, based on Mash Screen distance. *Reads* refers to the number of reads mapped to those genomes. *Coverage* refers to the amount of each genome that was covered by mapped reads. *Identity* refers to the average identity of the covered portions based on naive consensus from the pileups. *Score* and *P-Value* are results from Mash Screen.

| Organism | Reads | Coverage | Identity | Score | P-Value |
| --- | --- | --- | --- | --- | --- |
| *Propionibacterium acnes* HL072PA1 | 36,222 | 7.29% | 96.36% | 0.874 | 3.49e-136 |
| *Escherichia coli* strain 2014C-3250 | 59,744 | 4.99% | 95.16% | 0.837 | 7.32e-47 |
| *Proteiniclasticum ruminis* DSM 24773 | 751,538 | 76.57% | 91.83% | 0.930 | 0 |
| *Streptococcus parasanguinis* strain C1A | 74,807 | 57.50% | 95.84% | 0.942 | 0 |

### 2.2 What’s in my sequencing run?

As illustrated above, Mash Screen can discover unexpected organisms in sequence mixtures. This can be useful for checking any sequencing run for accidental contamination. While exploring containment results for a number of SRA datasets, we identified a number of potential contamination events. As an example, SRA run ERR024951 is a 1.1 Gb short-read dataset labelled as *Salmonella enterica* subsp. enterica serovar Typhimurium. However, analysis with Mash Screen also indicated the presence of *Klebsiella pneumoniae*, with strain k1037 (GCF_900086185.1) being the closest hit. We verified the presence of the contaminant by mapping all reads to both *Salmonella enterica* and *Klebsiella pneumoniae* reference genomes. This revealed that the majority of reads in this sample were in fact from *Klebsiella pneumoniae* (Table 2). The metadata on this particular dataset notes that it is part of a multiplexed sequencing run, suggestive of a possible contamination source. Because Mash Screen is extremely fast to run, it can be used as part of a standard sequencing workflow to catch such events before data is submitted to the public archives.

**Table 2.** Read mapping to confirm contamination of *Klebsiella pneumoniae* in a run, ERR024951, labelled as *Salmonella enterica*. *Organism* refers to either the labelled genome or the contaminant. *Reads* refers to the number of reads mapped to those genomes when both were provided as references in tandem. *Coverage* refers to the amount of each genome that was covered by the mapped reads. *Identity* refers to the average identity of the covered portions based on naive consensus from the pileups. *Score* and *P-Value* are results from Mash Screen.

| Organism | Reads | Coverage | Identity | Score | P-Value |
| --- | --- | --- | --- | --- | --- |
| *Salmonella enterica* subsp. *enterica* serovar Typhi str. CT18 | 1,962,751 | 88.91% | 98.64% | 1 | 0 |
| *Klebsiella pneumoniae* strain k1037 | 10,726,719 | 98.76% | 99.96% | 0.999 | 0 |

In the above examples, we show how Mash Screen can identify a ‘shortlist’ of genomes that are likely to be in a sequencing run. This can be generalized to the problem of metagenomic classification, where reference databases are often too large to allow traditional sequence analysis techniques such as read mapping. While not a classifier in its own right, Mash Screen can help address this problem by quickly filtering out reference genomes that are not well contained by the query mixture. This can drastically reduce the number of reference sequences necessary for good mapping and/or classification results. To test this application, we compared the known truth of the Shakya metagenome to the results of screening the reads for containment of RefSeq genomes. The filtered genomes amount to a significant size reduction in terms of the fraction of nucleotides represented, but still encompassed the vast majority of the known constituent genomes (Table 3). Note that a threshold of 1 (perfect containment) leaves many genomes out because Mash Screen is sensitive to both identity and completeness, and it has been observed that coverage of some genomes in this sample is low [17]. This experiment is thus a good illustration of the tradeoff between database size and classification sensitivity that may be necessary. However, in this case, pre-screening to a Mash containment of 0.9 retains all correct reference genomes while significantly reducing the database size, and tightening the threshold to 0.99 retains 56/58 of the correct genomes while reducing the database size an additional order of magnitude. Thus, Mash Screen can be helpful for recruiting a suitable set of reference genomes for metagenomic read mapping.

**Table 3.** Potential reduction in classification database size by pre-screening, illustrated using constituents of the Shakya metagenome. *Containment* refers to the threshold of the Mash containment score used to filter the genomes in the database, with 0 meaning no filtering and 0.9 meaning filter all genomes with a containment of less than 90%. *Positives* refers to how many of the known constituents had scores above that threshold, and thus would pass pre-screening (6 of the 64 constituents were not included because they have since been removed from RefSeq and thus were not in the reference database). *Database genomes* refers to the total number of genomes in the database with scores above the threshold, and *Database gigabases* refers to the total number of bases in those genomes.

| Containment (≥) | Positives | Positives (%) | Database genomes | Database gigabases |
| --- | --- | --- | --- | --- |
| 0 | 58/58 | 100% | 83,327 | 330 |
| 0.9 | 58/58 | 100% | 1,585 | 5.76 |
| 0.99 | 56/58 | 96% | 227 | 0.878 |
| 0.999 | 53/58 | 91% | 76 | 0.237 |
| 0.9999 | 43/58 | 74% | 58 | 0.182 |

### 2.3 Estimating proteome containment with a translated screen

Mash Screen introduces support for an amino acid alphabet, including 6-frame translation of the input reads. This allows for protein sketching and proteome containment estimation directly from metagenomic sequencing reads, which can be helpful when searching for divergent genes or genomes using short reads. To demonstrate that Mash’s k-mer-based approach can effectively measure protein similarity, we compared Mash Screen to the commonly used translated alignment program DIAMOND [19]. For a reference set, we began with reported protein fragments from two samples of the Human Microbiome Project [20], SRS015937 and SRS020263, and recruited their best hits to full-length proteins in the NCBI nr database, resulting in 28,334 protein sequences. Sequence reads from only one sample, SRS020263, were then mapped with DIAMOND to this combined set of proteins to ensure a broad range of alignment scores. Note that Mash and DIAMOND are designed for different tasks. Mash estimates global similarity, whereas DIAMOND computes local alignments. To estimate global similarity from DIAMOND alignments, a consensus residue was computed at each aligned reference position by simple majority, and the total number of matches was divided by the reference protein length to estimate global similarity, accounting for both similarity and coverage. By default, and as run, DIAMOND reports the single best HSP (i.e. mapping) for up to 25 reference sequences for each read. The consensus scores thus describe how well each protein is represented in the reads, without assigning each read exclusively to one protein, analogously to Mash Screen. The consensus scores were compared to the Mash containment scores computed by screening the sequencing reads against a sketch library consisting of the same reference proteins. Overall correlation between the two scores was *r*^2^= 0.64 and the relationship remained linear even for proteins with low coverage (Fig. 4). As was seen in the above comparisons to the Mash distance, the Mash containment score tended to overestimate the pairwise DIAMOND scores, especially with lower scores, but in a fairly consistent way.

**Figure 4.**
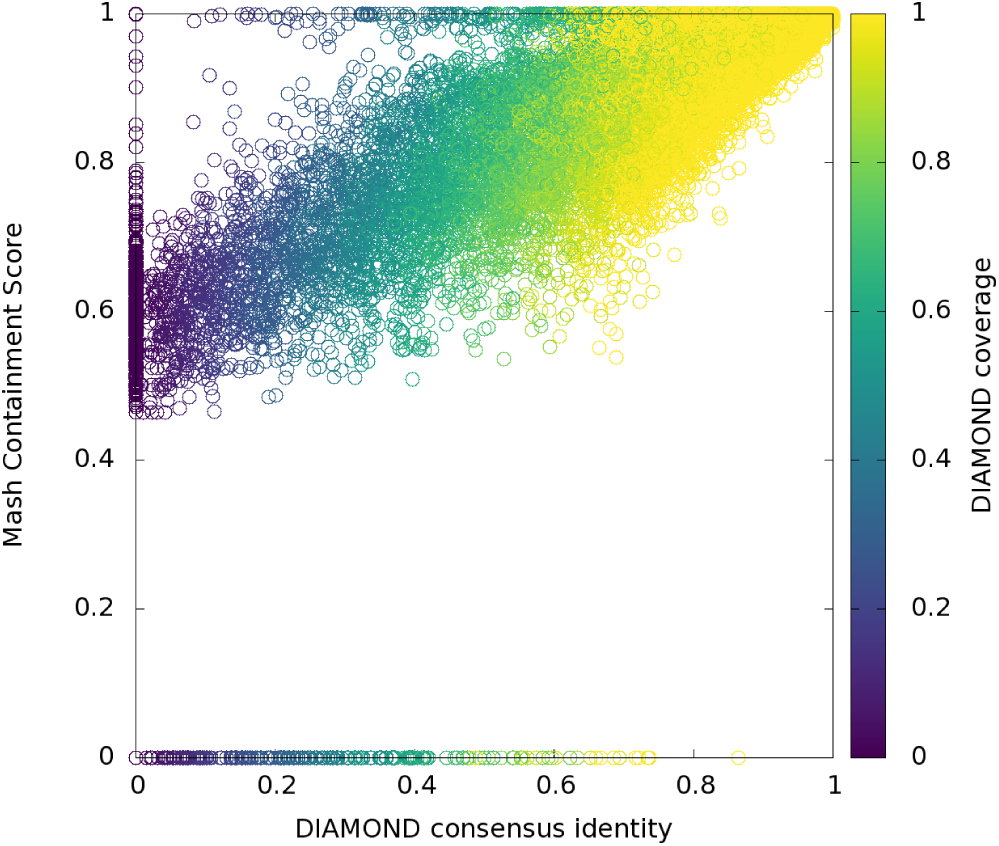
Correlation of Mash Screen containment scores with DIAMOND read mapping identity, each using six-frame translations to compare nucleotide reads to protein reference sequences. Each point represents a gene from the recruited set chosen for the experiment. Position on the *x*-axis corresponds to global alignment identity estimated from mapping, while position on the *y*-axis represents the containment score for the same protein as reported by Mash Screen. Coloring represents the fraction of the reference protein covered by aligned reads.

Sequence sketching confers its biggest advantage when reference sequences are long. Thus, in the case of individual proteins, Mash Screen is only slightly more efficient than a mapping-based tool such as DIAMOND (41 versus 121 CPU hours in the above experiment), because the resulting sketches are nearly as large as the original proteins. Instead, Mash Screen is designed for measuring the containment of whole genomes and proteomes within many metagenomes. Because screening a whole proteome sketch requires no additional space or runtime compared to screening a single protein, Mash scales extremely well in this case, whereas using a mapping-based tool would be infeasible. The below sections illustrate how this scalability can be used to process the entirety of the NCBI Sequence Read Archive to make new genomic discoveries.

### 2.4 Screening all metagenomes in the Sequence Read Archive

Mash Screen was designed to process the enormous collection of metagenomic data available in the public sequence archives, and scales linearly with the size of the input — requiring only slightly more time than would be required to simply read a FASTQ file. We leveraged this efficiency to process the entire metagenomic catalog of the NCBI Sequence Read Archive, and compute a containment score for every genome and plasmid in NCBI RefSeq versus every metagenome in the SRA. At the time of this analysis, there were 144,909 metagenomic sequencing runs (totalling ∼4 Pb) and 103,346 RefSeq genomes and plasmids (totalling 1.14 Tb) in NCBI. Note that during analysis, a small number of runs (between 1 and 2%) could not be accessed due to removal from the database, access restrictions, or repeated failed server requests (lists of the queried runs are provided in supplemental material). The resulting table, although sparse, represents a total of ∼15 billion containment estimates. Total runtime was ∼500,000 CPU hours (∼1,000,000 for six-frame translated), which is a feasible to recompute for each major RefSeq release with parallelization. Comparatively, appending additional SRA runs to the full table is trivial and requires a cost of just 8 CPU minutes per 1 Gbp of sequence (16 CPU minutes for six-frame).

Here, we make available the results of the RefSeq-SRA containment screen, consisting of all data available as of August 2018. To limit the size of these tables, we provide the results filtered in two ways: one table contains matches above a 0.95 containment threshold, while another relaxes this threshold to 0.80 but requires 3x median k-mer multiplicity. The former is useful for finding runs that contain a specific target, while the latter is useful for discovering novel sequences, as demonstrated in the following section. Screens were run using both untranslated nucleotide and translated protein screens, resulting in a total of four tables, available in supplementary materials (see Section 5.3).

### 2.5 Global distribution of human polyomavirus species in the SRA

Polyomaviruses are chronically shed from epithelial surfaces and are known to be common constituents of clinical and environmental samples. Although recent sero-epidemiological studies have confirmed that most humans are immunologically exposed human polyomaviruses HPyV 1 through 11, seroreactivity against HPyV12, New Jersey polyomavirus (NJPyV), and Lyon-IARC polyomavirus (LIPyV) is puzzlingly rare [21, 22]. It is thought that polyomaviruses generally co-speciate with their hosts and, accordingly, polyomavirus phylogeny tends to recapitulate the phylogenetic relationships of host animals [23]. Application of this principle led to the recent suggestion that HPyV12 may have been a laboratory contaminant that ultimately originated from shrew specimens [24]. Similarly, LIPyV was originally detected by PCR in human saliva but shows low seroreactivity in the general population [21], suggesting an external source for the original detection (e.g. household pets).

We selected all SRA metagenome runs with a proteome containment score ≥ 0.8, a median k-mer multiplicity ≥ 3, and a p-value ≤1 *×* 10^-100^ for at least one human-associated polyomavirus. Figure 5 shows containment scores for the 456 (of ∼143,000 total) runs meeting these criteria. The results show that most known human-tropic polyomaviruses were contained in multiple metagenomes (Figure 5). In contrast, HPyV12, NJPyV and LIPyV were not detected in any metagenomes. The result adds complementary support for serological and phylogenetic evidence that human infection with HPyV12, NJPyV, and LIPyV is rare or nonexistent.

**Figure 5.**
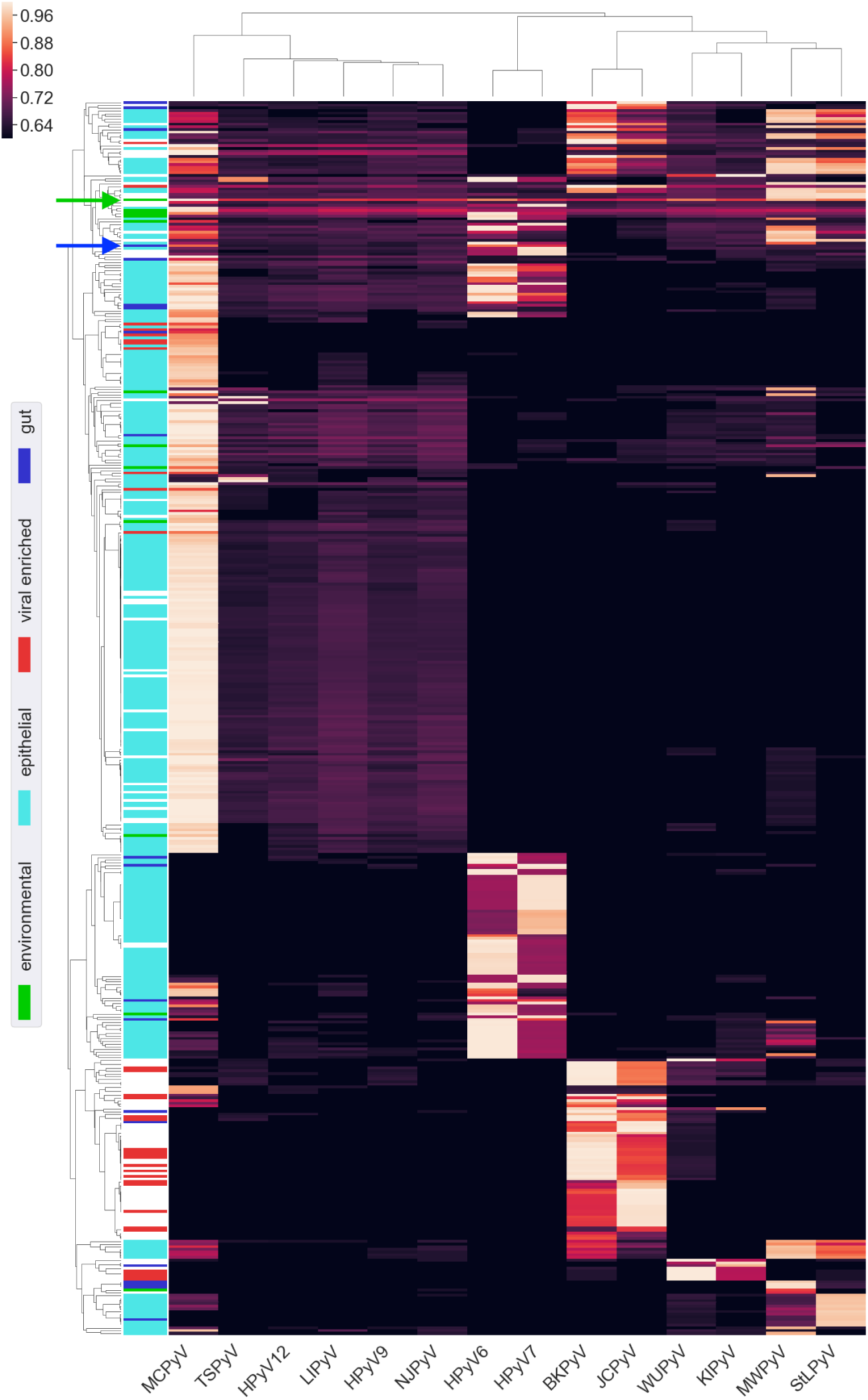
Metagenomic SRA runs with hits to human polyomaviruses. Hierarchical clustering was applied to both axes, and, where possible, descriptions from the *organism* metadata field were used to label various categories, depicted in the colored bands on the left. The vast majority of PyV-positive epithelial samples (cyan) are skin metagenomes from an eczema study (BioProject PRJNA46333). The blue arrow indicates the fecal sample in which QPyV was initially detected (SRR2565980). As an example environmental sample with strong hits to multiple human polyomaviruses, the green arrow indicates a money metagenome (SRR5256705).

NJPyV was initially detected in an immunosuppressed woman who was exposed to flood waters in the wake of Hurricane Sandy [25]. The virus was detected by in situ hybridization in myositic and cutaneous lesions, conclusively demonstrating productive infection of the patient. In contrast to HPyV12 and LIPyV, NJPyV shows close (70-80%) nucleotide similarity to polyomaviruses found in old world primates. A possible explanation for these observations is that NJPyV is a bona fide human-tropic virus but its prevalence is extremely low.

To search for additional examples of the hypothetical NJPyV category, we screened for SRA runs containing good (but not perfect) hits to known polyomavirus species. One identified run was a metagenomic analysis of human fecal samples from an 85-year-old hospital patient in Montreal, Canada (SRR2565980). The top polyomavirus hits for this run were HPyV6 and HPyV7, with k-mer multiplicities of 2 and 3, and proteome containment scores of 0.88 and 0.82, respectively. Lack of a containment score ≥ 0.90 to any known polyomavirus was interpreted as evidence for a potentially novel virus. All reads from this metagenome were assembled with metaSPAdes, revealing a nearly complete polyomavirus related to HPyV6 and HPyV7, which was manually finished to recover the complete genome. This new virus, for which we suggest the name Quebec polyomavirus (QPyV, GenBank BK010702), is 80% identical to HPyV7 and 67% identical to HPyV6 at the nucleotide level. As additional evidence for the new virus, another run (SRR2565981), representing an additional time point for the same patient, also had Mash containment scores passing the threshold (0.82 and 0.84, with median k-mer multiplicity 3 and 6, respectively, for HPyV6 and HPyV7). Investigation of a third time point from the same patient (SRR2565982) also revealed 4 reads mapping to the new QPyV assembly, which was below the Mash detection threshold. Although the high degree of similarity of QPyV to HPyV6 and HPyV7 strongly suggests that it is a primate-tropic polyomavirus, QPyV was only detected in fecal samples from a single study of hospital patients. It remains unclear whether QPyV (or NJPyV) should be thought of as a truly human-tropic polyomavirus.

## 3 Discussion

### 3.1 Limitations

Mash Screen quickly measures the containment of a set of reference sequences within a metagenomic mixture. Note, however, that each of the reference sequences is treated independently, without consideration for redundancy or overlap among others. These measurements do not, in themselves, estimate the composition of a metagenome but rather measure how well each reference genome is contained within the queried metagenome. As demonstrated above, these containment scores can be used to identify sample contamination, recruit a set of reference genomes for metagenomic analysis, establish the absence of specific reference genomes in public databases, and retrospectively discover novel genomes.

As with other k-mer based analyses, the containment estimate conflates sequence divergence with genome coverage. For example, a low-abundance species in a metagenome will typically receive a low containment score, simply because much of a query genome may not be fully represented by the sequencing reads. It may possible to disentangle these factors statistically using the multiplicity of the k-mers observed in the metagenome (e.g. as a proxy for sequence coverage). Conceptually, higher multiplicity for matching k-mers should indicate that k-mers not observed in the mixture are more likely to be missing due to sequence divergence, rather than low sequence coverage. Mash Screen currently tracks and reports the multiplicity of every hash in the reference sketches, but further work is needed to exploit this information fully.

Mash Screen is designed to efficiently estimate the containment of whole genomes or proteomes, and can also be used to detect the presence of any set of related sequences (e.g. plasmids, functional pathways). However, the efficiency of MinHash is lost when the reference sequences are individual genes and proteins. In these cases, the sketch of a gene can be as large as the gene itself. While Mash will operate normally under these conditions, searching for short sequences within metagenomes is better left to traditional sequence search and mapping tools.

### 3.2 Future directions

In addition to searching for contamination, we also demonstrated the potential of using a containment screen to pre-filter a metagenome classification database. This would be especially useful for tools that rely on mapping metagenomic sequence reads to a reference genome database. However, more rigorous validation of this use case is needed to prove its value. It may also be valuable to explore possible integration with classification pipelines to filter databases dynamically, given that database management is a burdensome step for many classification tools.

By definition of the MinHash process, similar reference genomes will share many minimum hashes between them. This makes the Mash Screen reference database highly compressible and keeps memory requirements low. In its current implementation, an off-the-shelf hash map is used for tracking reference sketch containment. As reference databases continue to grow, it may be desirable to improve the performance of this index. This could be achieved via a minimum perfect hash function for mapping k-mers to reference sketches, which would require only a few bits per k-mer to store the reference sketches [26].

### 3.3 Summary

Our online approach to containment estimation offers advantages over purely pre-computed approaches, namely that Mash sketches can now be used for both genome resemblance and containment operations. We validated Mash Screen by correlating pairwise resemblance with containment, and demonstrated its utility for contaminant screening and genome discovery. Further, we leveraged the efficiency of our method to compute the containment of all RefSeq genomes and plasmids within all SRA metagenomes, providing the research community with an unprecedented resource for retrospective data mining.

## 4 Methods

### 4.1 Mathematical foundations

Mash Screen compresses and indexes a database of reference sequences and measures their containment within a metagenome (the mixture). When estimating containment, all k-mers of the mixture are considered, ensuring adequate sensitivity. Because of this, any method for sampling the reference k-mer sets uniformly at random would suffice. However, for compatibility and to improve indexing efficiency, the reference sequences are sketched using the same bottom sketch strategy as Mash. Formally, a bottom sketch is defined as:

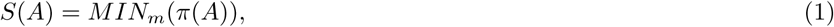

where *π*: Ω *→* Ω is a min-wise independent permutation [27] (typically approximated with a hash function) and *MIN*_*m*_(*W*) is the first *m* elements of the ordered set *W*. Computing bottom sketches for the reference sequences maintains compatibility between the resemblance and containment operations of Mash, as well as improves the efficiency of indexing. Because bottom sketching is a form locality sensitive hashing, the sketches of two similar sequences will contain many of the same hashes. This reduces the necessary index size for reference databases containing many similar sequences (e.g. strains of the same microbial species).

To estimate the containment of a set of reference k-mers *A* within some mixture *B*, we compare the sketch of *A* against all of *B*:

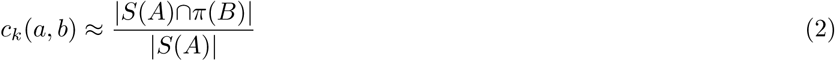

Since, for an arbitrary subset *A*^*′*^*⊆A*, every element of *A*^*′*^*∩B* must also be an element of *A∩B*, it follows that if *A*^*′*^ is chosen uniformly at random from *A, c*_*k*_(*a, b*) will be an unbiased estimate of the containment with expected error *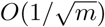*.

Lastly, it is convenient to convert the containment index into an estimate of sequence identity between the reference and mixture. Similar to the Jaccard index, the containment index drops exponentially with decreasing sequence identity. This is due to the fact that a single character substitution between two sequences affects *k* k-mers overlapping that position. Thus, to estimate identity from the containment index, we use a binomial model of k-mer survival. Given two sequences of equal length with a sequence identity of *i*, and assuming k-mers are unique and independent, the probability that any k-mer in *A* will be shared with *B* is *i*^*k*^. Because *c*_*k*_(*a, b*) represents the fraction of k-mers in *A* shared with *B*, we have:

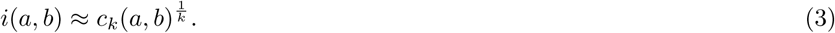

which we refer to as the Mash containment score.

#### 4.1.1 Amino acid estimation

Mash Screen supports six-frame translation of reads against reference sketches of protein sequences. This is performed by including hash values for all six translations of each amino acid k-mer in the MinHash containment estimation (c.f. 4.2.1). The out-of-frame elements are not expected to match *S*(*A*) and will not affect the containment score since *c*_*k*_(*a, b*) is independent of the size of *B*. While very large read sets may cause some erroneously-translated matches by chance, this is captured by a p-value, described below, that can be used to ensure only significant hits are reported.

#### 4.1.2 Assessing significance

In MinHash resemblance computations, erroneous k-mers (i.e. those arising from sequencing error) can crowd correct k-mers out of sketches, reducing sensitivity. This is not an issue when screening a read set, since, if present, the correct k-mers will be observed regardless of any other k-mers that may contain errors. It is, however, important to consider the effects of such k-mers on *specificity*, given that the hash of an erroneous k-mer could coincidentally match that of a k-mer in the reference. This is especially true when performing six-frame translation, given that, lacking *a priori* knowledge of the correct reading frame, five spurious k-mers must be emitted for every true one, even in the absence of sequencing error. We address this by formulating a p-value to estimate the significance of a distance estimate given the background of the observed k-mer space. Because the containment index does not depend on the size of the larger set, the expected containment of one random set, *X*, within another, *Y*, is simply the probability *r* that a random k-mer, *K*, appears in *Y*:

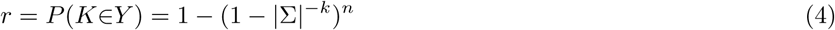

where *n* is the number of distinct k-mers in *Y*, Σ is the alphabet, and *k* is the length of each k-mer. Thus, the probability that *Y* matches *x* out of *s* k-mers in the sketch of *X* is given by the binomial distribution:

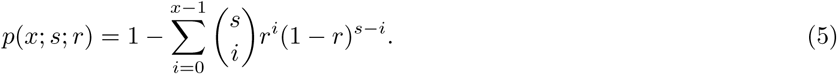

For six frame translation, the set size *n* is based on all amino acid k-mers emitted, ensuring that the risk of false positives from saturation of the k-mer space is reflected in the p-value. It should be noted that this value does not consider multiple testing, but can be scaled by the number of sketches in the reference set to attain an expectation.

### 4.2 Implementation

Mash Screen is integrated into Mash and included in all releases from v2.0 and higher. It is invoked as a subcommand, `screen`, similarly to `dist`, which is used computing the standard Mash distance. Unlike `dist`, however, `screen` must be given a pre-processed multi-sketch file to screen the reads against. This sketch file can be created with the `sketch` command in the same manner as for `dist`. If the sketch is created from protein sequences, the reads given to `screen` will automatically be translated into all six frames during screening.

#### 4.2.1 Streaming

As the mixture is intended to be an arbitrarily large set, such as a sequencing run, it is not desirable to load the entire set into memory. In contrast, the reference sketches are small and can easily be indexed in memory. To compute the containment of each reference sketch, the mixture is streamed against this index (Fig. 2). First, all distinct hashes present in the constituent sketches are collected into a reference set. This set is used as the keys to a map, where the associated values are counts for how many times the hash appears in the mixture, set initially to zero. Sequences from the mixture are then streamed, translating to six frames if applicable, and all their k-mers hashed, incrementing the counters in the map if a key for that hash is present. This process is parallelized by using atomic operations to increment to the counters without locking the map. Finally, a distance for each constituent is determined by tallying how many hashes in its sketch have non-zero counts in the map. Note that the number of entries in this map, and thus memory used, will depend on the number of distinct hashes in the complete set of possible constituents. Since many reference genomes will be highly similar to one another, storing a MinHash sketch for each reference is more efficient than storing a random selection of k-mers (as noted above).

#### 4.2.2. P-value

The p-value (described in Eq. 5), used to estimate the significance of a given distance estimate, relies on the number of distinct k-mers |*B*| observed in the streamed reference set. Counting this number exactly would require an entry for each of these k-mers in memory, erasing the memory efficiency afforded by the streaming estimation algorithm. Instead, this count is estimated from the maximum hash value of an auxiliary MinHash sketch of the reference set, constructed as the set is streamed for this purpose [28]. If the sketch size is *s*, its maximum hash value is *v*, and each hash uses *b* bits, the number of distinct k-mers in the genome is estimated as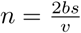.

### 4.3 Experiments

#### 4.3.1 Correlation between Mash distance and containment estimates

Simulated reads were generated using the `art_illumina` command with parameters`-f 50` (50-fold coverage), `-l 100` (100-base read length), and `-ss HS20` (Illumina HiSeq 2000 platform profile), and with the combined 64 reference genomes as input. Mash was run with defaults for both `screen` and `dist`, except the use of `-p` for multiple threads. RefSeq genomes were binned to Shakya reference genomes with a custom script.

#### 4.3.2 Correlation of Mash Screen containment scores with DIAMOND read mapping identity

Blast was run with default parameters (using the `blastp` command), a custom script was used to collect identifiers of best hits, and `blastdbcmd` was used to extract these sequences from `nr`. The `diamond` subcommands `makedb` and `blastx` were run using default parameters, with the exception of parameters to control parallelism and output formats. The SamTools [29] `mpileup` command was used to create pileups from the DIAMOND mappings, and a custom script was used to determine consensus identity and coverage from the pileups.

#### 4.3.3 Screening all metagenomes in the Sequence Read Archive

SRA accessions for runs labeled as metagenomic were extracted with Bioconductor [30]. Reads for each run were streamed to an instance of `mash screen` using `fastq-dump` from the SRA Toolkit [4] with the `-Z` flag. Default options were used for `mash screen` except `-p 2` to specify 2 threads. Results for each run were filtered and compiled into tables with custom scripts.

#### 4.3.4 Novel virus assembly

The SRA project SRP064400 was downloaded using `fastq-dump` from the SRA Toolkit [4], and assembled using metaspades [31] with default parameters. Assembled contigs were screened using `blastx` and a custom blast database containing a select set of conserved proteins from DNA viruses [32]. Contigs containing likely viral protein sequences were further screened all known viruses using `blastn`. Contigs with less than 90% nucleotide identity to any known viral genome were then manually annotated and analyzed for completeness, which revealed a nearly complete genome for a new polyomavirus related to HPyV6 and HPyV7. The viral genome was completed using methods previously described by Pastrana and colleagues [32].

## 5 Declarations

### 5.1 Ethics approval and consent to participate

Not applicable.

### 5.2 Consent for publication

Not applicable.

### 5.3 Availability of data and materials

The datasets generated and/or analyzed during the current study are available at https://mash.readthedocs.io/en/latest/data.html.

### 5.6 Authors’ contributions

The screening algorithm was designed by BDO and AMP, and implemented by BDO. Validation using the Shakya metagenome was performed by BDO. SRA experiments were performed by BDO and AK. MinHash protein space experiments were performed by BDO and AS. Viral discovery, assembly, and analysis was performed by GJS, CBB, AK, and AMP. Classification database reduction experiments were performed by BDO and SK. The study was conceived and coordinated by AMP. All authors read and approved of the final text.

## 5.7 Acknowledgments

We thank Brian Walenz for helpful discussions about implementation details. This research utilized the computational resources of the NIH HPC Biowulf cluster (https://hpc.nih.gov).

